# Live-cell imaging of pathogenic fungal hyphae reveal dynamic cellular responses to clinical antifungals

**DOI:** 10.1101/2024.07.19.602466

**Authors:** D.D. Thomson, R. Inman, E. M. Bignell

**Affiliations:** Manchester Fungal Infection Group, Division of Infection, Immunity and Respiratory Medicine, Faculty of Biology, Medicine and Health, University of Manchester, CTF Building, Grafton Street, Manchester M13 9NT, UK; Medical Research Council Centre for Medical Mycology at the University of Exeter University of Exeter, Geoffrey Pope Building Stocker Road, Exeter, EX4 4QD, UK

## Abstract

Antifungal susceptibility testing quantifies end-point fungal biomass in liquid cultures initiated from non-invasive yeast or spore morphologies. However, end-point analyses obscure informative spatio-temporal responses to drug exposures. In the major fungal pathogens *Aspergillus fumigatus* and *Candida albicans* we used microfluidic-coupled, fluorescence-mediated live-cell imaging to capture the real-time responses of fungal hyphae to clinical concentrations of AmBisome or Caspofungin. In both fungi, AmBisome exposure caused rapid growth arrest, extensive hyphal vacuolation and membrane blebbing. Responses to Caspofungin exposure were slower with initial lytic effects occurring after 1.5 or 4 hours in *A.fumigatus* and *C.albicans*, respectively. Whilst *C.albicans* hyphae undergo unsalvageable hyphal lysis in response to Caspofungin, *A.fumigatus* exhibit several compensatory growth behaviours, including a novel resuscitative growth form, that circumvent lytic events to maintain apical and sub-apical hyphal growth. This study reveals how the differing biologies of the two pathogens affected outcomes and contributes to the highly disparate rates of antifungal efficacy amongst commonly used drugs, where spore/yeast-derived inhibitory doses may underestimate the dose required to arrest/kill the invasive hyphal morphotypes of fungal pathogens *in vitro*.

## Introduction

Microdilution broth testing is the primary method utilised for routine determinations of minimum inhibitory and effective concentrations (MICs and MECs, respectively) of antifungal drugs [1, 2]. Antimicrobial Susceptibility Testing (AST) protocols involve multi-well static liquid culture of fungal yeast or spore suspensions in the presence of antifungal agents to detect, via visual inspections and optical density measurements, the minimum drug concentrations that impede fungal growth. Although simple to implement and highly reproducible in the clinic, AST analyses employ end-point growth determinations of the pre-invasive (non-hyphal) fungal morphologies exposed to drug. Therefore, the effect of drug exposure on hyphal growth, the invasive form of the fungus most commonly found in patients, are missed. Such oversights may have important consequences for clinical management of fungal infection, for example, it has been reported that differing morphologies of the same fungal isolate exhibit different MICs [3].

Longitudinal analyses of antifungal treatments to pathogenic fungi have focused largely upon the action of the cell wall-active echinocandin, Caspofungin. Caspofungin induces hyper-branching and tip lysis followed by de novo intra-hyphal growth within the lysed hyphal tips when initially applied to *A.fumigatus* spores at clinically-relevant doses [4-6], or hyphae at significantly-higher [7] concentrations. Similarly, *C.albicans* yeast cells subjected to half-MIC Caspofungin on solid media also generated hyphae susceptible to tip lysis [8] as well as exhibiting a modified cell wall ultrastructure and lipid localisation after MIC drug exposures [9-12]. These powerful studies have revealed intriguing cellular responses to the echinocandin drug class, identifying novel and shared antifungal resistance strategies amongst fungal pathogens, such as active cell wall remodelling and morphological aberrancies. They also identify a need to align routine testing practices with clinically relevant culture conditions with renewed focus on the invasive morphotypes expected at infection sites. Moreover, such studies highlight the dearth of information on the dynamic modes of action of other antifungal drugs. For example, despite decades of approved clinical use to treat fungal infections, the dynamic cellular responses to liposomal Amphotericin B (AmBisome) have yet to be documented. AmBisome is a lipid formulation of Amphotericin B, which specifically delivers Amphotericin B to fungi to extract ergosterol from the fungal cell membrane like a sponge, resulting in fungicidal activity against both yeast and hyphae [13, 14]. Only in *Saccharomyces Cerevisiae* yeast has ergosterol extraction been found to be 80% complete in <30 mins under Amphotericin B, quickly followed by cell death [14].

Microfluidic live-cell imaging (Mf-LCI) in multi-wells is a powerful tool for observing and comparing real-time responses of living fungal cells to antifungal drugs [7]. In contrast to AST end-point analyses, Mf-LCI permits the longitudinal and comparative analyses of dynamic responses of invasive hyphal cells to perfused drug exposure, thereby better emulating the clinical scenario of established invasive disease. Furthermore, the in-built flexibility of drug delivery regimens permits real-time analysis of fungal responses to introduction, withdrawal and reintroduction of antifungal drugs as well as user-determined emphasis on specific fungal morphotypes. We applied Mf-LCI to perform a comparative analysis of synchronously captured responses of living *A.fumigatus* and *C.albicans* hyphae, to Caspofungin and AmBisome exposure. This study describes observed distinct and shared cellular adaptations, occurring over differing timescales, in two major human fungal pathogens in response to clinical antifungals.

## Results & Discussion

To follow the responses of actively growing fungal hyphae to antifungal drug exposure we used Mf-LCI and fungal strains expressing genetically encoded fluorescent reporters to visualise and quantify fungal biomass, cellular integrity (lysis or leakage), vacuolation and growth. In order to study clinically relevant *in vitro* responses of fungi, the EUCAST-determined [1, 2] inhibitory drug concentrations were determined for each pathogen with Caspofungin and AmBisome (Supplemental Table 1).

### *A.fumigatus* antifungal response

Upon perfusion with 0.5 µg/ml (MEC) Caspofungin, pre-germinated *A.fumigatus* hyphae continue to extend in an apparently normal manner until apical lysis occurred at 75 mins post-exposure (Movie 1 hyphae a and b 03:15). The remaining hyphae in exemplar Movie 1 lysed within 105 mins of each other (75-180 mins). As previously described [6, 7], frequent and heterogeneous hyphal outcomes were observed, including apical and subapical compartment lysis (Movie 1 hypha c 03:45 & 05:00), subapical branching (Movie 1 hypha c 04:00-08:00) and *de novo* intra-hyphal growth (Movie 1 hypha c apex 04:45-08:00). Intrahyphal apical growth of a new hyphal tip originates from the apical septum and continues within the lysed apical compartment, past the site of previous tip lysis [6, 7]. In contrast we additionally observed an example of retrograde intra-hyphal growth upon lysis of a basal hyphal compartment. Here, *de novo* intra-hyphal growth occurred from the local branch-septum back into the lysed basal compartment containing the mother spore (Movie 1 hypha c spore compartment; Movie S1). Immature germlings lacking septa (i.e. non-compartmentalised) were unable to display any form of residual growth once lysed and no longer exhibited fluorescence indicating a complete loss of metabolic activity after 6 hrs Caspofungin perfusion (data not shown; [15]).

We observed a novel type of regenerative growth behaviour (designated here as resuscitative growth) involving full recovery of intracellular cytosolic fluorescence and growth capacity; without the need for *de novo* hyphal growth that typifies the intra-hyphal growth response [6]. Resuscitated hyphae took between 15 min (Movie S2; hypha a) and 75 mins (Movie 1 hypha a; Movie S3) to presumably repair lysed cell wall, recover turgor and cytosol, which collectively facilitated apical resuscitated growth after Caspofungin-induced tip lysis. Resuscitated tips were still susceptible to subsequent lytic events. Although this phenomenon was observed without the presence of extracellular features (Movie S2; hypha a), in the majority of instances, resuscitative growth was associated with the prior appearance of a highly localised post-lytic extracellular fluorescent signal at the apex of lysed hyphae (Movie 1 hypha a; Movie S3). Amongst an exemplar population of 12 lysing hyphal tips (Movie 1), 4 hyphae exhibited the post-lytic extracellular apical fluorescence signal. Three of these hyphae underwent the novel resuscitative hyphal growth programme involving replenishment of fluorescence and turgor within the lysed compartment (Movie 1 hypha a). A fourth lysed hypha exhibiting the residual extracellular apical fluorescence signal did not resuscitate, but instead continued growth by branching from its basal septa, out-with the lysed apical compartment (Movie 1 hypha b; Movie S1 hypha b). Subapical hyphal compartments also illustrated post-lytic resuscitative growth (Movie 1 hypha c 03:45) which were then capable of septum formation (2 hrs later), indicating it is not a phenomenon exclusive to hyphal tip recovery. The post-lytic cellular responses we map here in time, occurring at MEC drug concentrations, may contribute to the failure of perfused Caspofungin to sterilise *A.fumigatus* hyphae after 48 hrs *in vitro* (data not shown) and possibly *in vivo*.

In stark contrast to Caspofungin, *A.fumigatus* hyphae perfused in the same manner with 0.125 µg/ml AmBisome elicited an immediate and homogenous cessation of growth (<15 mins; Movie 1 far right), which did not recover after 6 hours perfusion. This rapid response to MIC AmBisome was associated with subsequent vacuolation (Movie 1 far right), where the peak number of vacuoles were formed in hyphae after approximately 45 mins of AmBisome exposure (Movie 1 far right).

As growth was completely inhibited by MIC AmBisome and vacuolation progressed, amongst an exemplar population of 24 fungal hyphae, three subsequent responses were observed: 11/24 hyphae maintained their vacuoles (46 %; Movie; hypha 1d), 8/24 hyphae underwent vacuole lysis after 1hr 45 mins exposure (33 %; Movie 1; hypha e) and 5/24 hyphae underwent cellular lysis after 4 hrs 15 mins exposure (21 %;Movie 1; hypha f). Vacuole lysis was followed by extinction of cytosolic fluorescent signal in one hypha within the subsequent 2 hrs (Movie 1; hypha e; 05:30-07:30) suggesting a possible leaching of cytosol, since photobleaching was not observed in other hyphae within the studied timeframe (Movie 1; site g). AmBisome treated hyphae were incapable of any growth after perfusion with fresh medium without AmBisome (data not shown).

### *C.albicans* antifungal response

Similar to observations of *A.fumigatus*, we found that exposure of *C.albicans* hyphae to MIC Caspofungin (0.08 µg/ml) did not completely inhibit hyphal growth after 6 hrs of perfusion (Movie 2). Hyphae continued growing and did not begin lysing until 4 hrs of perfusion in the Mf-LCI platform (Movie 2; site a; 08:15).

Unlike *A.fumigatus, C.albicans* hyphae do not exhibit apical filamentous salvage growth behaviours following tip lysis. Instead, we observed hyper-branching (Movie 2; site b), followed by mass hyphal tip lysis and switching to isotropic budding growth (Movie 2; 08:00-10:00). Although the time to begin lytic activity was more than double in *C.albicans* (4-8 hrs) compared to *A.fumigatus* (1-2hrs), the period to achieve lytic activity amongst the original hyphae was similar between the two pathogens under MIC/MEC Caspofungin (2 hrs). A longer-term population-switch to budding growth at branch sites (Movie 2; site b; from 07:15) and leading hyphal tips (Movie 2; site c; 07:45-10:00) then dominated the growth response. By 6 hrs exposure to perfused Caspofungin, newly formed isotropic cells were capable of continued budding growth, albeit still susceptible to lysis. Our observations indicate that *C.albicans* yeast derived from lysed hyphae can maintain a steady-state level of residual growth during MIC Caspofungin treatment *in vitro* under EUCAST-compliant Mf-LCI. We additionally observed that if the *C.albicans* microcolony was dense prior to Caspofungin perfusion and hyphal lysis, budding growth was maintained and somewhat protected from lysis in the dense central mass (Movie S4), possibly representing a drug penetration defect. Further, we found that reintroduction of drug-free RPMI-MOPS media after 6 hrs of Caspofungin exposure at 37 °C resulted in no recovery to filamentous growth. Instead, further steady-state budding growth and lysis of fluorescent yeast cells continued for 5 hrs in the absence of drug (Movie S5).

Similar to *A.fumigatus*, in response to AmBisome MIC treatment (Movie 2), *C.albicans* hyphae were observed to immediately form vacuoles (<15 mins) and cease growth. These vacuoles subsequently lyse (Movie 2; beginning at 08:15), leading to diminishing cytosol fluorescence in hyphae (Movie 2; site d;), possibly due to the fungal membrane becoming porous via AmBisome’s action on membrane ergosterol [13, 14]. Suspected fungal membrane detachment was observed under carefully tuned differential interference contrast (DIC) optics, where we visualised lipid-like protrusions originating along the hyphal wall after 2hrs 15 mins of perfusion with MIC AmBisome (Movie 2e). These were more pronounced and balloon-like after 45 mins of perfusion with 4 µg/ml AmBisome (Movie S6). This is the first known live-cell visualisation of a membrane ‘blebbing’ phenomenon in fungal cells, a process that is visually akin to mammalian cell blebbing where cytosolic leakage occurs via localised regions of compromised cell membrane. Growth arrest in AmBisome-treated *C.albicans* cells was similar to *A.fumigatus* in a highly homogenous manner and time-frame, this may be due to the ubiquitous non-localised distribution of the drug target (ergosterol) along the fungal hyphae. The observed en-mass vacuolation in hyphae presumably reflects attempts to sequester imported Amphotericin B, as seen in *C.albicans* studies of drug uptake using fluorescence lifetime imaging [16], and may be associated with an active reduction in cytosolic volume or an arrest in the G1 cell cycle [17, 18].

To track and quantify fungal 2D cross-sectional area, in response to drug perfusion, we applied 4D image processing to Mf-LCI data over multiple biological replicates in the presence or absence of drug for both pathogens (Figure 3). MEC Caspofungin treatment of *A. fumigatus* hyphae resulted in continued growth, at a rate comparable to that observed in absence of drug, for approximately 2 hrs. A subsequent drop in relative growth rate was consistently observed for all hyphae analysed, and can be attributed to hyphal lytic events described earlier. Fungal growth did subsequently continue at a linear but reduced rate after mass lytic events; the net effect of multiple salvage growth behaviours described from exemplar Movie 1.

**Figure 1.**
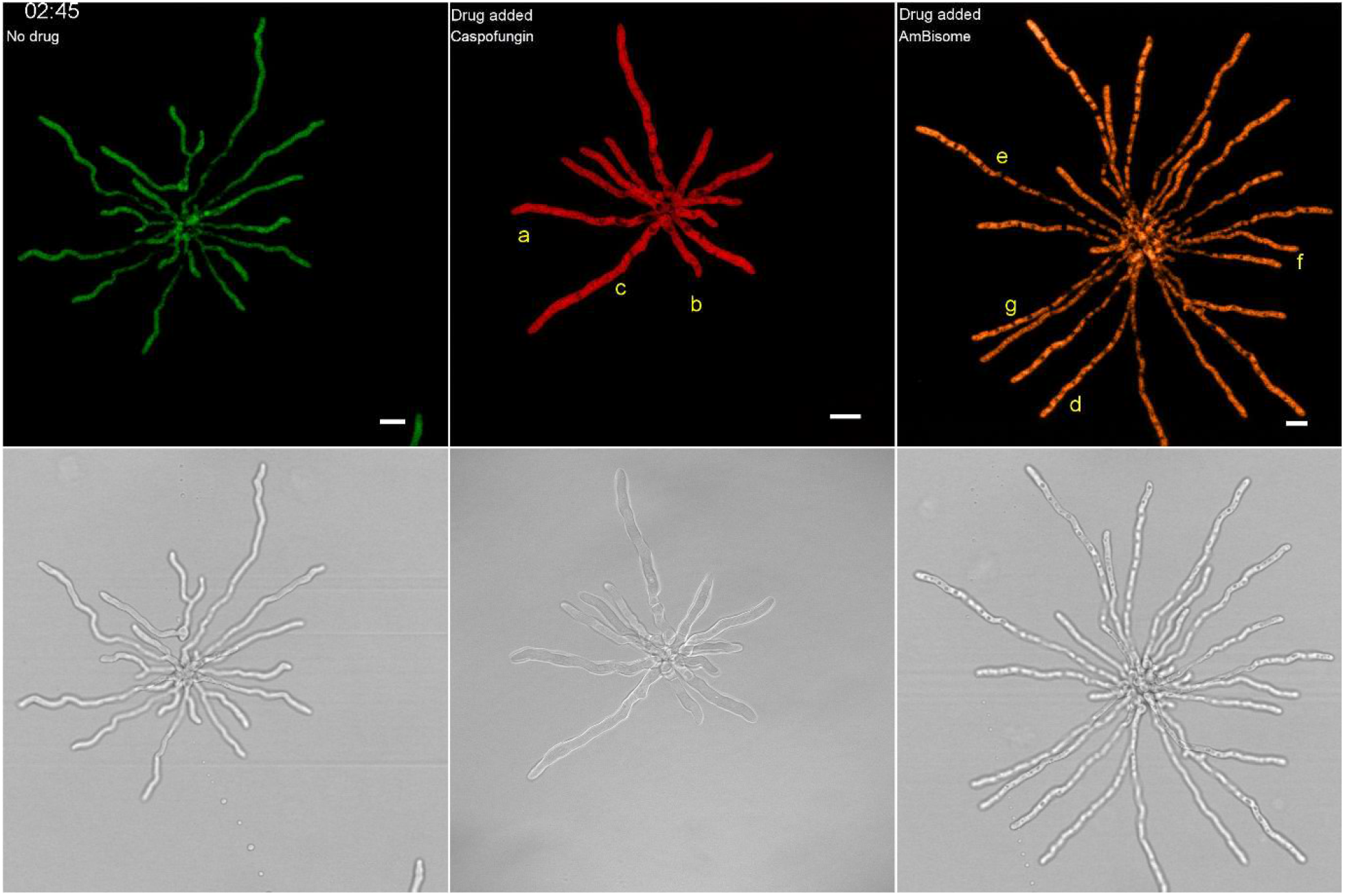
Montage from Movie 1 of fluorescent *A.fumigatus* responding to 0.5 µg/ml Caspofungin and 0.125 µg/ml AmBisome. Untreated (green; left column), Caspofungin (red; middle column) and AmBisome (amber; right column) treated *A.fumigatus* hyphae growing in a Cellasic microfluidic chamber. After 2 hours of growth in media without drug, new media (green) or drug (red/amber) were perfused into the chamber. The fluorescence images were pseudo coloured in the top row, while the bottom row displays the transmitted light images. Hyphae denoted with the following letters indicate key events occurring in Movie 1:***a) Caspofungin treated hyphal tips lyse, maintain an extracellular fluorescent tip signal and resuscitate. Subapical branching is also evident in the compartment behind the apex.***b) Caspofungin treated hyphal tips lyse, maintain a fluorescent apical tip and do not resuscitate. Subapical branching is evident in the basal compartment.***C)Caspofungin treated hyphal tips lyse and undergo intra-hyphal growth at the nearest apical septa. The subapical compartment also undergoes resuscitative growth with septum formation. The basal compartment undergoes retrograde intra-hyphal growth from the basal branch septum toward the mother spore body.***d-g)AmBisome treated hyphae undergo vacuolation followed by: d) maintenance of vacuoles e), vacuole lysis and diminishing cytosolic fluorescence protein, f) sudden hyphal lysis, and g) vacuole lysis with no diminishing of cytosolic fluorescence protein. Scale bar = 10 µm.

**Figure 2.**
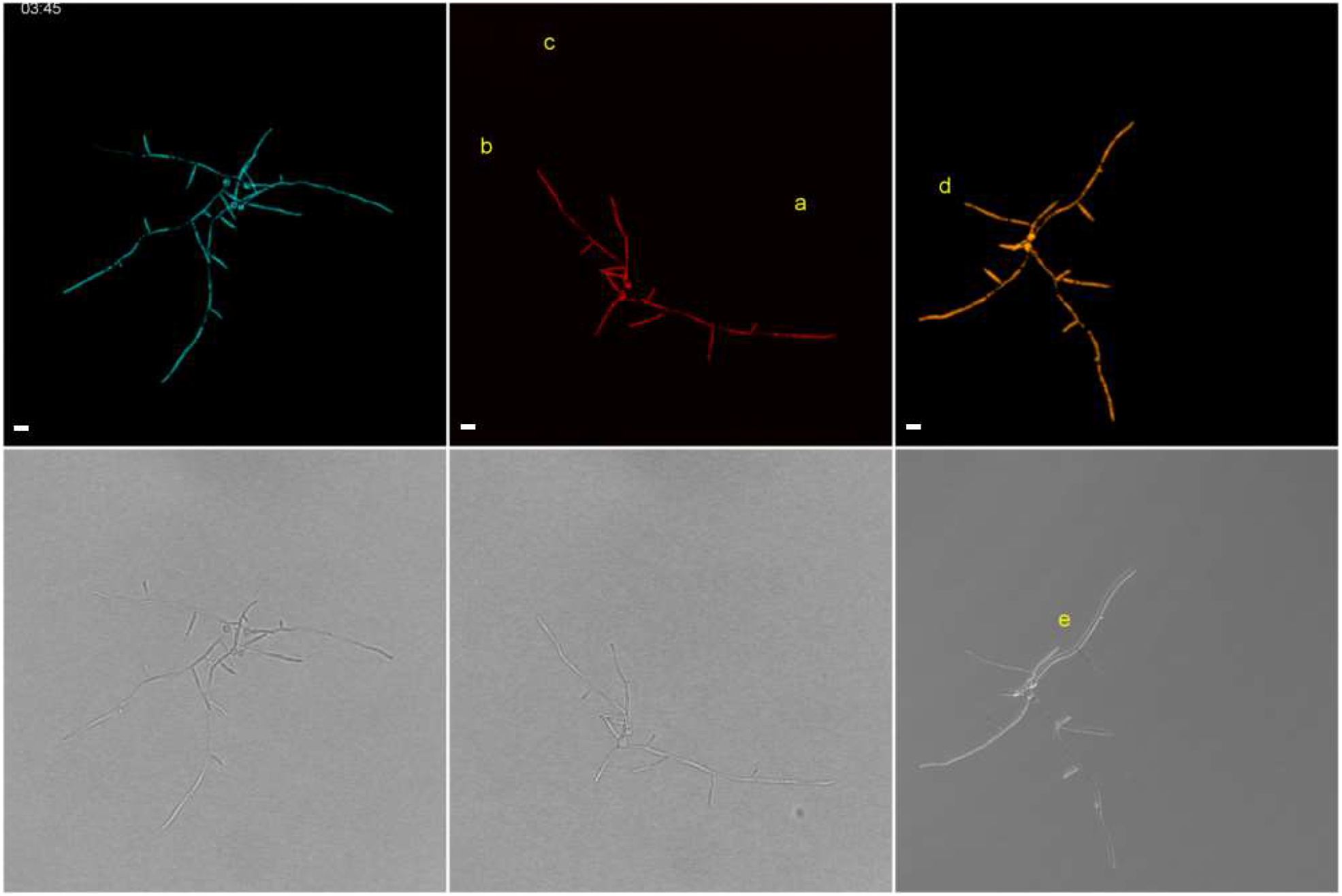
Montage from Movie 2 of synchronised movies of fluorescent *C.albicans* responding to 0.08 µg/ml Caspofungin and 0.5 µg/ml AmBisome. Untreated (cyan; left), Caspofungin (red; middle) and AmBisome (amber; right) treated *C.albicans* hyphae growing in a Cellasic microfluidic chamber. After 4 hours of normal growth media, new media (green) or drug (red/amber) are perfused into the chamber. The fluorescence images were pseudo coloured in the top row, while the bottom row displays the transmitted light images. Hyphae denoted with the following letters indicate key events occurring in the movies:***a) Caspofungin treated hyphae begin lysing.***b) Caspofungin treated hyphae begin branching and becomes less polarised and reverts to budding growth.***c) Caspofungin treated hyphal tips begin septating and reverting to budding growth.***d) AmBisome treated hypha undergoes vacuolation followed by vacuole lysis and a diminishing of fluorescence protein.***e) AmBisome treated hypha undergoes fungal cell blebbing. Scale bar = 10 µm.

**Figure 3.**
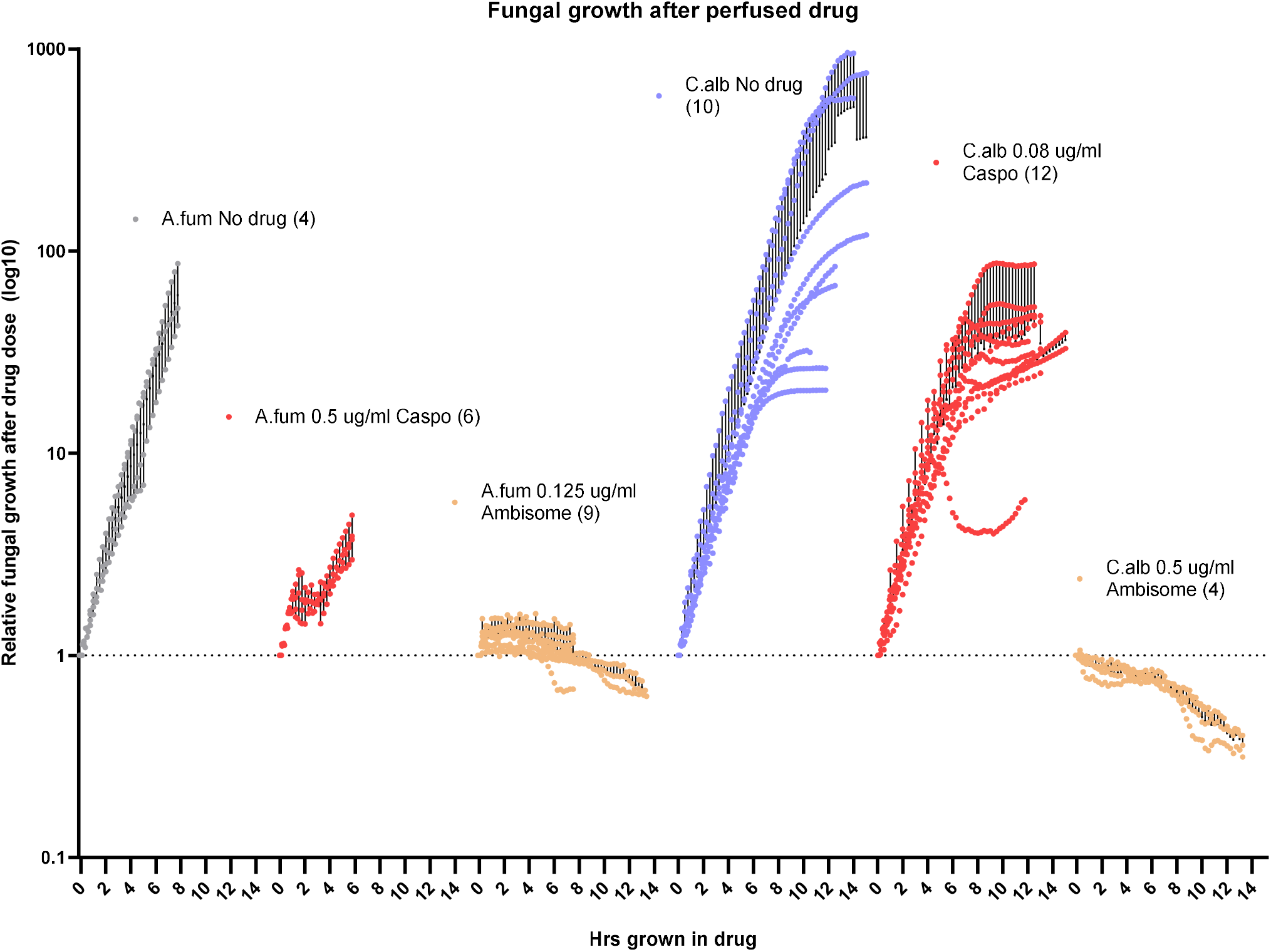
Image-based fungal growth dynamics to antifungal agents, based on segmented fluorescent cytosol images. Each data point is expressed as relative growth upon drug perfusion. Black bars represent the standard deviations to the mean. Biological replicates are indicated in brackets of data legend.

The growth plots display very dynamic growth responses to Caspofungin which differed between pathogens, whereby MIC-treated *C.albicans* hyphae continued growth at a comparable rate to untreated hyphae for 4 hrs to 8 hrs exposure, depending on the starting size of the *C.albicans* mass upon drug treatment. The reduction in linear relative growth is attributed to the synchronous en masse hyphal tip lysis events within each of the *C.albicans* micro colonies. Unlike *A. fumigatus*, where hyphal lysis events occurred within a consistent time of exposure (2-4 hrs), the timing of hyphal lysis between individual *C.albicans* microcolonies was more sporadic (4-8 hrs). The plateau regions of the relative *C.albicans* growth data reflect the switch to budding growth, where the majority of newly produced yeast are also lysing, creating a steady-state equilibrium of fungal mass.

Both pathogens consistently shared the dramatic response of immediate growth cessation (< 15 mins) and presumed loss of fungal cytosol, upon MIC AmBisome exposure. Unlike *C.albicans*, we did not observe fungal membrane blebbing in *A.fumigatus* hyphae.

## Conclusions

In this study we undertook a comparative analysis of the dynamic hyphal responses to Caspofungin and AmBisome exposure. We address how hyphae, rather than the more-often studied yeasts or spores, respond to continuous perfusion of antifungal drugs under EUCAST-guided growth conditions. Indeed, via modified MIC testing, higher concentrations of drug were required to inhibit growth of *A.fumigatus* germlings, compared to spores (Table S1), a phenomenon previously described for AmBisome [3]. These data support the view that spore-derived inhibitory doses may underestimate the inhibitory doses required to arrest/kill the subsequently formed invasive morphotypes of fungal pathogens *in vitro*.

Synchronised Mf-LCI demonstrates *in vitro* that, under continuous Caspofungin perfusion, hyphae of the filamentous mould *A. fumigatus* remain viable and maintain hyphal growth via distinct modes (branching, hyper septation, intra-hyphal growth and the newly-described resuscitative growth). A new phenomenon of residual post-lytic apical cytosolic aggregates (the underlying basis of which is not currently understood) were identified to frequently associate with hyphal resuscitation, possibly via the localised repair of ruptured cell wall. Despite the ability to rejuvenate via such modes of growth, hyphae remained susceptible to Caspofungin-induced lysis. In the time-frame analysed, fungal biomass relative to untreated hyphae was drastically reduced 10-20 fold. In contrast to the delayed growth inhibition to Caspofungin, AmBisome immediately stopped all growth within 15 mins of exposure and only lost mass over time, via loss of fluorescent protein from lysed or possibly porous cells.

In response to MIC Caspofungin, the hyphae of the polymorphic fungal pathogen *C.albicans* lyse and switch to budding yeast-form growth, with varying dynamics that appear to be dependent upon colony size. The *C.albicans* switch to isotropic growth under Caspofungin may have clinical implications for biofilm integrity and/or systemic dissemination in the blood stream. *C.albicans* similarly stopped growing and underwent vacuolation and fluorescence cytosol loss in response to MIC AmBisome.

This study describes real-time responses of fungal hyphae to antifungal drugs, documenting for the first time under clinically relevant antifungal drug exposure, the occurrence and dynamics of highly prevalent fungal lysis and novel re-growth events *in vitro*. It will be important to understand these events in a clinical context, where the immune system may be engaged by lysed and/or polymorphic fungal material evoked by antifungal treatment regimens. These responses may contribute to emergence of echinocandin resistance seen in the clinic and better-inform future combinatorial treatment regimens to abate echinocandin-driven resurgent growth.

These findings present new forms of resuscitative antifungal growth, which occur alongside classical antifungal responses, and clarifies the real-time hyphal responses to MIC AmBisome in two major fungal pathogens.

## Methods

### Fungal strains and culture conditions

*A.fumigatus* ATCC 46645 expressing cytosolic YFP [PgpdA::yfp(ptrA)] spores were streaked on Potato Dextrose Agar plates, cultured for 48 hrs at 37 °C and harvested in water via Miracloth filtration, prior to Mf-LCI. The yeGFP *C.albicans* strain (CAI4) *RPS1/RPS1*::*pACT1-GFP* [19] was a gift from Alistair Brown (University of Exeter), streaked on Yeast Peptone Dextrose(YPD) agar plates at 30 °C for 48 hrs. Colonies of *C.albicans* yeGFP yeast were picked from YPD agar plates and inoculated into YPD liquid culture for overnight incubation at 30 °C and diluted in RPMI-MOPS prior to Mf-LCI.

RPMI-MOPS (pH 7) fungal growth media were prepared by following the EUCAST guidelines [1, 2]. Working concentrations of AmBisome were prepared for each experiment by adding 12ml dd H2O to a vial of Amisome powder to generate a vial of 4 mg/ml. This was used to generate a final concentration of Amphotericin B (within the AmBisome formulation) in RPMI-MOPS as indicated in the results text. Stock Caspofungin was prepared to 10 mg/ml in DMSO. Working concentrations of Caspofungin were prepared in dd H2O for final dilution in RPMI-MOPS as indicated in the text.

Hyphal MIC testing of *A.fumigatus* was performed by germinating 1-2.5×10^5^ spores/ml in 100 µl RPMI-MOPS (pH 7) at room temperature overnight in a flat-bottomed 96 well plate, followed by an additional 8 hr incubation at 37 °C, until approximately 95 % of the spores had germinates and contained 1 septa. The plates were sealed with breath-easy sealing membrane to prevent evaporation. This was followed by the addition of 100 µl of RPMI-MOPS or RPMI-MOPS containing double strength antifungal to each well. Growth inhibition was then recorded by visual inspection after 24 and 48 hrs incubation at 37 °C.

### Microfluidic Live-cell imaging

Microfluidic cellasic plates were aspirated and perfused with RPMI-MOPS in all wells to expel the plate antibiotic. Fluorescent fungal cells were diluted to a concentration of 1×10^5^ cells/ml in RPMI-MOPS, where 200 µL of suspension was loaded into a YO4E Cellasic plate (Merck, UK). Fungal cells were perfused into the plate chambers via microfluidic bursts at a pressure of 8 PSI for 5 seconds using the ONIX2 software (v 5.0; Merck, UK). This was repeated until a satisfactory cell distribution was achieved. Fungal spores/yeast were left to adhere for 30 mins, whereby RPMI-MOPS was flowed into the chamber at 2 PSI continually to remove unadhered cells. Adhered cells were continually perfused with RPMI-MOPS until hyphae were established and ready for imaging. *A.fumigatus* spores were pre-germinated in the plate for 8-10hrs at 37 °C in RPMI-MOPS prior to live-cell imaging. *C.albicans* yeast were germinated in the microfluidic plate at 37 °C for 2-4 hrs, prior to live cell imaging. The plate was mounted on the microscope and a solution switching program was established via the ONIX Cellasic controller and software. A continual flow of RPMI-MOPS media was set for 2-6 hrs at 1 PSI, depending on the size of the microcolony, then followed by AmBisome or Caspofungin-containing RPMI-MOPS at initially 4 PSI for 5 mins, then 1 PSI, up to 14 hrs.

### Image acquisition

Under microfluidic perfusion, 3D confocal and widefield image stacks (7 slices, 3 µm apart) were acquired every 15 mins on a 37 °C temperature controlled (Cube & Box, CH) Leica SP8x confocal microscope (Leica Microsystems, Germany) equipped with a dry Leica 40x/0.85NA HCX PL APO CS, or, when required for the visualisation of fungal blebbing, an oil immersion 63x/1.4NA HC PL APO CS2 objective lens with tunable DIC optics.

Fluorescence images of yeGFP *C.albicans* and eYFP *A.fumigatus* strains were acquired with 488 nm and 514 nm argon lasers, respectively, with appropriate HyD point detection windows in LASX software (v.5.1.0 Leica Microsystems, Germany). The laser intensity and exposure to the cells were kept to a minimum (<1%) to avoid photobleaching and cellular phototoxicity. Associated transmitted images were acquired with a transmitted light detector. Additional widefield acquisitions were performed on a temperature regulated Zeiss AxioObserver Z1 equipped with a Hamamatsu Fusion sCMOS camera and a 20x/0.8NA Plan-apo lens with a GFP filter set (Zeiss #62) which captured both GFP and YFP expressing fungi.

### Image processing

Fluorescence 4D image data sets were imported into FIJI [20] and 3D projected by standard deviation. Time-series images were then registered in FIJI using StackReg [21] to eliminate spatial drift. Movie montages were then created with the *Combine* function and annotations were generated using the *Time stamper* and *label* features in FIJI. The cross-sectional area of fungal cytosol was quantified in fluorescence time-series images in FIJI for each culture condition. Briefly, each image was background subtracted and blurred with a Median filter prior to applying a Percentile threshold and water-shed of the fungal segmentations. This segmentation cross-sectional areas were then analysed using the *Particle Analyser* function in FIJI to report the total cross-sectional area of fungi per time point. Total fungal cytosol areas were then exported into Graphpad Prism and transformed into relative growth, after the point of drug perfusion, for each condition. Where there was no drug, the same time point of drug entry for other conditions was taken, as the initial relative mass.

## Acknowledgements

We wish to thank Alistair Brown for kindly providing the *Candida albicans* yeGFP strain and Sven Kraapmann for providing the *Aspergillus fumigatus* eYFP strain. We thank Sergio Moreno-Velasquez, the Mycology Reference Centre Manchester and Alexandra Brand for their valuable discussions. We also thank Ian Leaves for the technical assistance in reviving *C. albicans* strains from stocks.

This work was funded by Gilead Sciences and the Medical Research Council Centre for Medical Mycology at the University of Exeter (MR/N006364/2 and MR/V033417/1) and MRC project grants MR/M02010X/1, MR/S001824/1 and MR/L000822/1 and the BBSRC project grant BB/V017004/1 to EMB. This research was carried ou.t at the National Institute for Health and Care Research (NIHR) Exeter Biomedical Research Centre (BRC).

## Disclaimer

Movies 1 & 2 were commissioned and funded by Gilead Sciences and are under the copyright of Gilead Sciences, Inc. All rights reserved. Videos are used under licence from Gilead Sciences, who have had no input into the content of this research article.

EB received funds for speaking at symposia organized on behalf of Gilead. DT has received funds for consultation from Owlstone Medical in the last 5 years.

## Movies & supplemental movies

Each movie is embedded in the following Zenodo link to download: https://zenodo.org/records/12685210. If you require the original highest resolution version of these movies please contact the authors and we can make them available to you.

**Supplemental Table 1.** MIC data.

| Organism | Morphology | AmBisome (µg/ml) |  | Caspofungin (µg/ml) |  |
| --- | --- | --- | --- | --- | --- |
|  |  | MIC<sub>50</sub> | MIC<sub>90</sub> | MEC |  |
|  |  |  |  | MIC<sub>50</sub> | MIC<sub>90</sub> |
| *A.fumigatus*<br>ATCC 46645 | Conidial | 0.06 | 0.125 | 0.5 |  |
|  | Hyphal | 0.125 | 0.25-1 | 1 |  |
| *A.fumigatus*<br>ATCC 46645<br>YFP | Conidial | 0.06 | 0.125 | 0.5 |  |
|  | Hyphal | 0.125 | 0.25-1 | 1 |  |
| *C.albicans*<br>ACT1p-yeGFP | Yeast | 0.125 | 0.5 | 0.06-0.125 | 0.25 |

**Supplemental Movie 1**

***A.fumigatus* hyphae undergo retrograde intrahyphal growth under MEC Caspofungin**. Zoomed in movie of Hypha ‘c’ from Movie 1 undergoing mother-spore compartment lysis and subsequent retrograde intrahyphal growth into the lysed spore compartment. Left panel is the DIC frames of the movie with yellow arrows indicating retrograde growth from the branch septum into the lysed spore compartment of the hypha. Right panel indicates the cytosolic fluorescence disappearing upon lysis and new intrahyphal growth into the spore compartment.

**Supplemental Movie 2**

**Resuscitative growth occurs without extracellular apical mass**. Fluorescent *A.fumigatus* hyphae treated with MEC Caspofungin. Left panel contains the cytosolic fluorescence and the right panel is the associated DIC image. Hypha ‘a’ loses fluorescence upon apical lysis and undergoes resuscitative growth within 15-30 mins of lysis. Note that the bottom hypha ‘b’ undergoes resuscitative growth after producing a post-lytic extracellular apical mass.

**Supplemental Movie 3**

**Post-lytic extracellular apical masses may contain cytosol**. Fluorescent *A.fumigatus* hyphae treated with MEC Caspofungin. Left panel contains the cytosolic fluorescence and the right panel is the associated DIC image. The yellow arrow tracks the optically dense extracellular mass in the DIC frames as it loses fluorescence. The extracellular mass loses optical density and fluorescence within 45 mins.

**Supplemental Movie 4**

**Caspofungin treated hyphae can form dense microcolonies**. Fluorescent *C.albicans* hypha treated with MIC Caspofungin for 12 hrs, resulting in dense microcolony formation. Top panel is the cytosolic fluorescence, and the lower panel is the corresponding DIC images. Scale bar = 10 µm.

**Supplemental Movie 5**

**Washing Caspofungin out does not recover filamentous hyphal growth**. Yeast-reverted *C.albicans* already treated with MIC Caspofungin for 7 hrs, was solution switched to wash out the drug with RPMI-MOPS did not result in a return to filamentous hyphal growth. Top panel is the cytosolic fluorescence and the lower panel is the corresponding brightfield images. Scale bar = 10 µm.

**Supplemental Movie 6**

**MIC AmBisome treated *C.albicans* hyphae form extracellular bed-like membrane structures**. DIC timelapse movie show regions where balloon-like refractive structures originate and develop from a point in the fungal hyphal wall, indicated by white arrows. Time format is in hh:mm. Scale bar = 10 µm.

## Notes

https://zenodo.org/records/12685210

